# MICROBIOTA DRIVES THE SEXUALLY DIMORPHIC INFECTION OUTCOMES IN MEALWORM BEETLES

**DOI:** 10.1101/2024.09.10.611485

**Authors:** Srijan Seal, Devashish Kumar, Pavankumar Thunga, Pawan Khangar, Manisha Gupta, Dipendra Nath Basu, Rhitoban Raychoudhury, Imroze Khan

**Affiliations:** Trivedi School of Biosciences, Ashoka University, Sonepat, Haryana-131029; Indian Institute of Science Education and Research, Mohali (IISER-Mohali), Manauli, Punjab-140306

**Keywords:** Sexual dimorphism, Infection, Immunity, Microbiome, *Tenebrio molitor*

## Abstract

Sexually dimorphic responses to pathogenic infections in animals may stem from sex-specific differences in their life history and immune investment. Recent evidence highlights that such sex-specific variations in immune responses can also be critically regulated by microbiota. However, direct experiments to test how microbiota jointly impacts sex-specific immunity and vulnerability to pathogens are still limited. To this end, we used *Tenebrio molitor* beetles to first establish that sexes appear to differ in their microbiota composition and infection responses. Females were more vulnerable to bacterial infections and carried a higher bacterial load than males. When we depleted the microbiome, only females improved their post-infection survival, leading to a loss of sex-specific infection outcomes. Males, on the other hand, remained unaffected. Microbiota reconstitution (via feeding on faecal matter) of microbiota-depleted females increased their susceptibility to infection again, restoring the sexual dimorphism. We thus found a causal association between microbiome and infection responses. We also found reduced expression of an antimicrobial peptide tenecin 1 in females, which could be associated with their higher infection susceptibility, but such immune gene-vs-phenotypic associations were not consistent across microbiota manipulations. Immune strategies that are required to mediate the causal links between microbiome and infection response might thus vary with microbiota manipulations, warranting future investigations.

## INTRODUCTION

Pathogens are widespread (1), with considerable divergence in how they impact the infection outcomes across sexes (2). For example, females in multiple taxa, ranging from insects to birds and mammals, show lower post-infection mortality costs. Life history and sexual selection theories predict that such sex-dependent variations might arise from sexually dimorphic immune investments (3). For example, males experiencing more significant variation in reproductive success and intra-sexual competition might invest less in immunity, making them more susceptible to pathogens. In contrast, females evolving under strong selection maintain a long, healthy lifespan to maximise lifetime reproductive fitness (4), invest more in immunity, and are less susceptible to infections. However, there are many exceptions, too, where these sex-dimorphic patterns of immunity and infection outcomes can be reversed. They can either be pathogen-specific even within a single species (5) or influenced by pathogens which already adapted differently to each sex (6) or, sensitive to environmental variables (7) and sex-specific physiological changes (8).

Recent studies have highlighted that inherent divergence in microbiota also drives sexual dimorphism in immunity (9), which may have large implications for natural variability in infection outcomes. In male mice, a higher abundance of *Eubacterium* and *Clostridium* species in the gut boosted T-cell activity (10). Germ-free male mice had lower antibody levels than controls, but no such difference was found in females (11). Moreover, transplanting gut bacteria from male or female mice to germ-free mice of the same or opposite sex demonstrated sex-specific immune responses (11). While female recipients of the male microbiota showed the upregulation of immunoglobulin variants, recipients of the female microbiome had activated proteases expressed in mucosal mast cells. Also, in a study on fall armyworms *Spodoptera frugiperda*, eliminating gut bacteria led to downregulating Toll and Imd pathways in adult males but not females (12). Together, these findings thus conclusively prove that microbiota plays a pivotal role in regulating sex-specific immune responses. However, experiments that investigate the direct link between microbiota, immunity, and sex-specific infection outcomes in a unified experimental setting are limited.

We thus jointly analysed the role of the gut microbiome in maintaining sexually dimorphic immunity and infection outcomes in mealworm beetles *Tenebrio molitor*, infected with an entomopathogen *Bacillus thuringiensis*. Adult females were highly susceptible to infection, carried higher pathogen burden and showed lower expression of antimicrobial peptides than males. However, such strong sexually dimorphic infection outcomes and immunity completely disappeared when we depleted the beetle microbiota. Reconstitution of the microbial community could rescue the sexually dimorphic infection outcome, although underlying immune mechanisms may differ from untreated control beetles. Our results thus provide strong evidence for microbiota and immune responses interacting intimately to influence sex-specific differences in vulnerabilities to pathogenic infections.

## METHODS

### Beetle maintenance and microbiota manipulations

We performed our experiments with a laboratory-adapted population of *Tenebrio molitor* beetles reared at 30°C, with a generation cycle of 10-12 weeks (See **Fig. 1** for a brief experimental outline). We used virgin adult beetles (13–15 days old post-eclosion) to avoid any confounding effects of mating by collecting and segregating pupae into males and females with *ad libitum* food till the start of the experiments. Since wheat bran acts as their source of food and the environment they reside in (13), we altered the microbiome of freshly-eclosed beetles by maintaining them on different diets.

**Figure 1:**
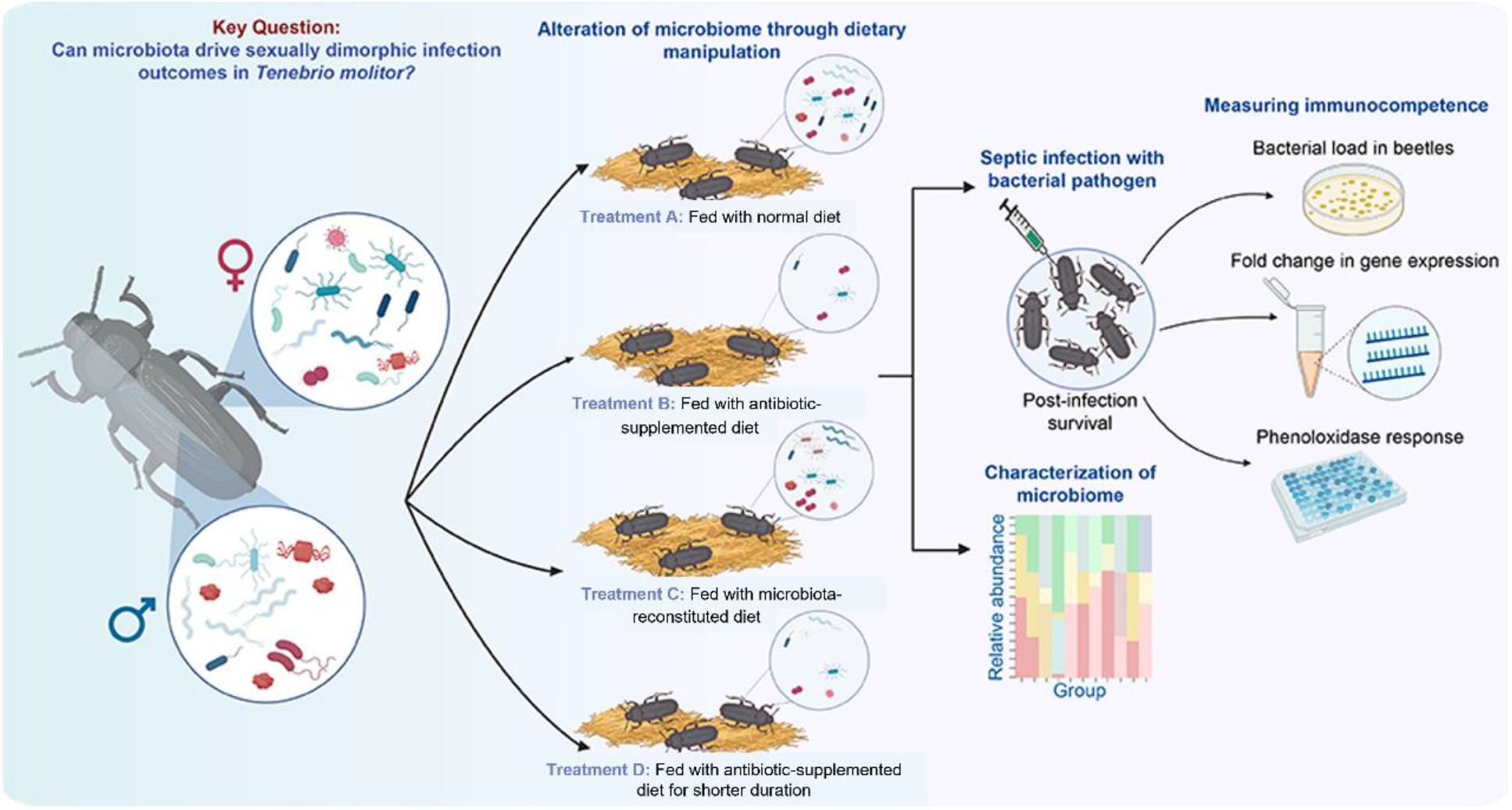
A brief outline of experimental design. Freshly eclosed adult beetles were either reared in (**Treatment A**) a normal diet (i.e., regular wheat bran), (**Treatment B**) an antibiotic-supplemented diet, (**Treatment C**) a diet supplemented with faecal matter to reconstitute the microbiota after initial depletion by feeding on antibiotic-supplemented diet for 6 days. Separately, we also included another treatment where beetles were reared in antibiotic-supplemented wheat bran for the first six days, followed by another 6-days in sterile wheat bran (i.e., shorter antibiotic exposure & no microbiota reconstitution; Treatment D). Subsequently, we performed a correlated estimate of their post-infection survival, pathogen clearance, and immune responses and characterised the changes in gut microbiome composition across sexes.

We maintained a set of beetles that were reared in normal wheat bran until day six post-eclosion but were then transferred to a sterile diet (i.e., autoclaved wheat bran), serving as procedural controls for the microbiota manipulations (**Treatment A**). Treatment A produced results similar to those of beetles that lived on a normal diet (i.e. regular wheat bran) until they were assayed (See **Fig 2A and 2B**) and, hence, can also serve as a proxy for unhandled full control beetles. We depleted the beetle microbiome by feeding them autoclaved wheat bran (i.e., sterile) mixed with the powdered form of two broad-spectrum antibiotics (Ampicillin and Streptomycin in 0.05% w/w ratio) for 12 days. We then transferred them to fresh autoclaved wheat bran for two days before infecting them (total 14 days) (**Treatment B**). Since both antibiotics have a half-life of less than 3-4 hours in animals, the gap of two days between antibiotic exposure and infection removed the confounding effect of any trace amount of antibiotics that might interfere with the infecting pathogen (14, 15). To restore the gut bacteria, we allowed beetles to feed on a diet supplemented with faecal matter (16). To this end, they were initially given antibiotics for six days, followed by another six days of exposure to autoclaved wheat bran, which had been pre-conditioned by 100 adult beetles and 100 larvae for a week (i.e., beetles excreted in the autoclaved wheat bran to restore microbiota) (**Treatment C**).

**Figure 2:**
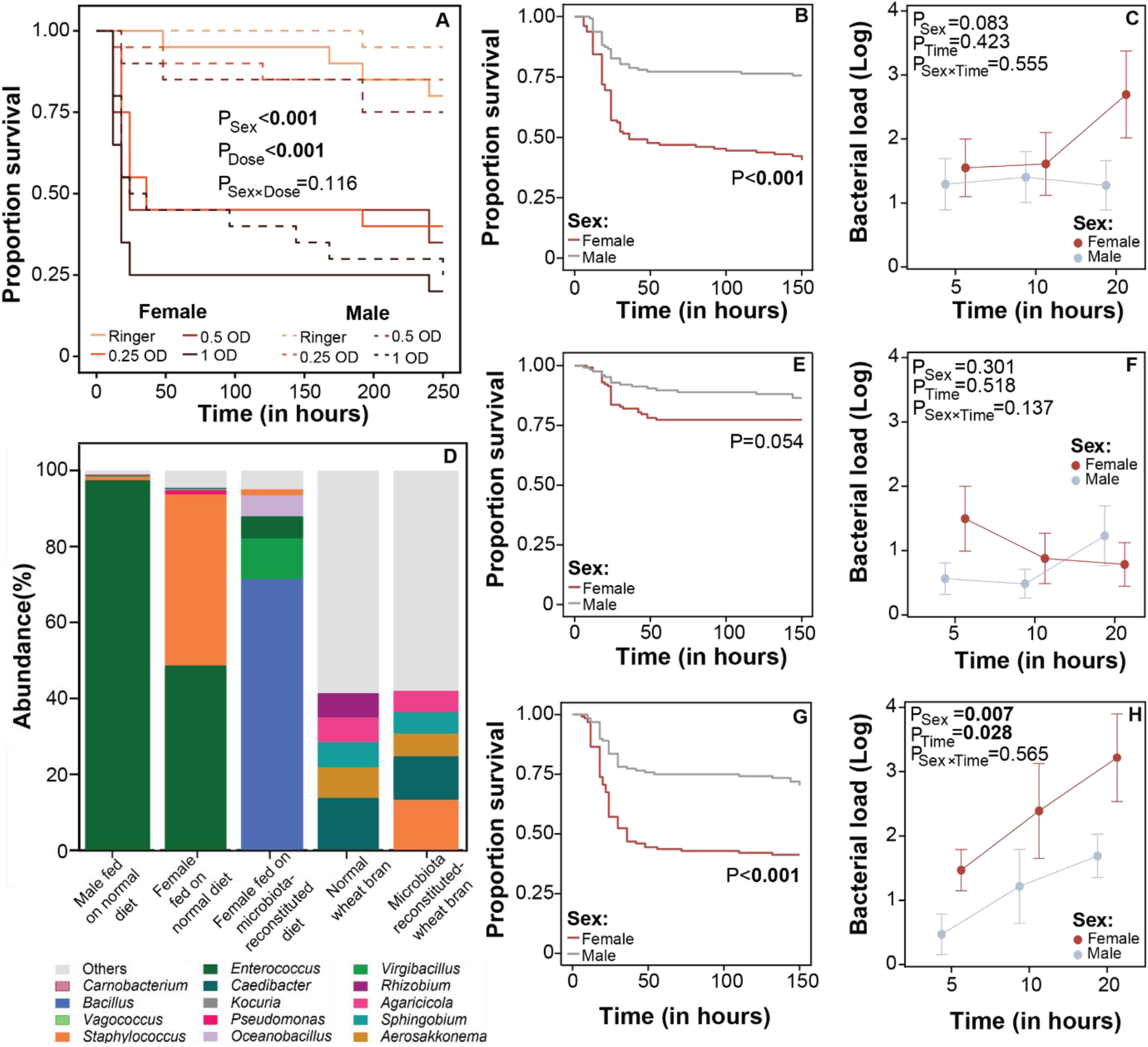
**(A)** Proportion of beetles surviving after infection with different doses (i.e., equivalent to 0.25, 0.5 and 1 OD_600_; or insect Ringer solution as mock infection) of *B. thuringiensis* (Bt) (n= 20/sex/infection dose). The P-value represents individual and interaction effects of sex and infection dose (data analysed using the Cox proportional hazard model); (**B**) Proportion of beetles surviving after infection with 0.25 OD (i.e., 720 ± 50 cells/beetle) of Bt when fed on a normal diet. **(C)** The corresponding bacterial load dynamics till 20hpi (n=10 beetles/sex/timepoint/ dietary treatment). **(D)** The microbiome present in beetles fed on a normal diet, females fed on microbiota-reconstituted diet, normal wheat bran and microbiota-reconstituted wheat bran. Stacked bar plots showing the percent abundance of the top 5 OTUs in each sample and the remaining OTU’s are grouped under “others” category (n=3 beetles pooled together/ sex/ dietary treatment). Post-infection survival of beetles infected with 0.25 OD Bt after feeding on **(E)** an antibiotic-supplemented diet and **(G)** microbiota-reconstituted diet (n= 24–32/sex/ dietary treatment/replicated trial). Bacterial load in beetles maintained under **(F)** an antibiotic-supplemented diet or **(H)** a microbiota-reconstituted diet (n= 10/sex/time point/dietary treatment). In panels B, E and G, P-values represent the effects of sex on post-infection survival (data analysed using mixed effects Cox proportional model). In panels C, F and H, P-values represent the main effects of sex, time-points and their interaction on log-transformed bacterial load (data analysed using a generalised linear model fitted to a Gamma distribution). Significant P-values are highlighted in bold.

Separately, we had another subset of beetles where they fed on antibiotic-supplemented bran for six days, followed by exposure to only autoclaved wheat bran (without antibiotics and microbiota reconstitution) to test the effects of exposure time to antibiotics and not reconstituting the microbiota (**Treatment D**). Similar to treatment A, other treatments also lasted 12 days, followed by two days of rearing under a sterile diet before beetles were infected and assayed. We provided beetles with cotton balls soaked in apple juice (with or without antibiotics based on the treatment) during this entire process. We replenished the antibiotic-supplemented food every alternate day.

To confirm the changes in beetle gut microbiome maintained under different treatments, we dissected the gut of uninfected individuals (males and females fed on a normal diet, antibiotic-supplemented and the reconstituted diet) on the 12th day (n=3 individuals/ dietary treatment/sex) and isolated microbial DNA using a Wizard genomic DNA isolation kit (Promega) following the manufacturer’s protocol. We also isolated microbial DNA from the normal and microbiota-reconstituted wheat bran following a similar protocol. We amplified the DNA using universal bacteria-specific primers (See **Table S1**). The resulting PCR products were pooled in equimolar concentrations for nanopore library preparation using the SQK-LSK 108 ligation sequencing kit (Oxford Nanopore technologies). Unique barcodes were added by ligating barcode adaptors using the EXP-PBC001 PCR barcoding kit (Oxford Nanopore Technologies). Subsequently, the barcoded samples were pooled in equimolar concentrations, sequencing adaptors were added, and the samples were sequenced on the MinION Flow cell (FLO-MIN106) R9 version (17) (See supplementary information, henceforth SI). We performed demultiplexing using the Guppy program (Oxford Nanopore Technologies) followed by filtration and trimming to obtain high-quality reads. Finally, we used the Lastal program in LAST v973 to identify bacterial communities (18). Separately, we also sequenced the samples from each beetle to understand the individual variations (See SI). However, we did not use any statistical test as the replicate size was low.

### Assays

We subjected the experimental beetles to the following assays (Detailed protocol is described in the SI)—

#### A. Post-infection survival

We began by checking the sex-specific post-infection survival with varying doses of Gram-positive bacteria *Bacillus thuringiensis* DSM 2046 (Bt) (19) or a Gram-negative bacteria *Pseudomonas entomophila* (Pe) (20) infection in beetles maintained under normal dietary conditions (See SI for details on infection protocol and dose). This enabled us to test the general sex-specific outcome of the bacterial infection. We conducted all analyses in R version 4.3.3. We used Cox proportional hazard analysis to test the main effects of sex and dose (wherever applicable) on post-infection survival using the “survival” package in R (21).

Subsequently, for microbiome-manipulated experiments, we only used a single dose of Bt cells (0.25 OD which is equivalent to 720 ± 50 cells/beetle) for the infection that induced ∼50% mortality in females. We repeated the experiment four times (n=24—32 beetles/sex/dietary treatment/trial). We analysed post-infection survival for dietary manipulation experiments with a Cox proportional mixed effects model with sex as a fixed factor and repeated trials as a random factor.

#### B. Bacterial load

We quantified the *B. thuringiensis* load at three-time points (5 hrs, 10 hrs and 20 hrs) across the infection window following a previously published protocol (22) (also see SI) (n= 10 beetles/sex/timepoint/dietary treatment). We analysed the bacterial load data using a generalised linear model fitted to a gamma distribution, with sex and time as fixed effects.

#### C. Immune responses

We measured the gene expression levels of two representative beetle antimicrobial peptides (AMPs), tenecin 1 and tenecin 4 (23), in infected (or mock-infected) males and females at 8 hours post-infection, that are likely responsive to Gram-positive and Gram-negative bacteria respectively (24) (n=7 beetles/sex/infection treatment). Within each sex, we calculated the relative fold changes in expression level between infected and sham-infected individuals, using the formula 2^-ΔΔCt^ as described in Schmittgen et al. (25), where ΔC_t_ represents the difference in C_t_ value between the candidate AMP gene and rpl27a (ribosomal protein subunit used as internal reference) (See **Table S2**). We also tested the sex-specific gene expression level of tenecin 1 with both microbiota depletion (treatment B) and reconstitution (treatment C). We analysed the fold changes in relative gene expression between the sexes using a generalised linear model fitted to the Gamma distribution.

Parallelly, we measured the changes in phenoloxidase activity upon infection across sexes following a previously published protocol (13) in beetles fed on a normal diet (treatment A), antibiotic-supplemented diet (treatment B), and microbiota-reconstituted diet (treatment C) (n=10 beetles/sex/infection treatment). We analysed the impact of sex on the phenoloxidase activity of beetles fed on different dietary treatments using the Wilcoxon rank sum test.

## RESULTS

### Females were more susceptible to bacterial infection and carried a higher pathogen burden

We found that females were more susceptible to both *B. thuringiensis* (**Figure 2A, B Table S3)** and *P. entomophila* infection (**Figure S1, Table S4**) than males under standard dietary conditions.

Subsequently, when tested with increasing doses of Bt cells, females continued to show higher mortality than males, regardless of the infection dose. Subsequently, we found that *B. thuringiensis* cells grew more in females (**Figure 2C, Table S5**) but not in males. Males could restrict bacterial growth consistently at lower levels during the first 20 hours post-infection, suggesting that males were more resistant to bacterial infection.

### Microbiota drove the sex-specific effects of bacterial infection

We identified patterns of notable differences in gut microbiota composition between females and males. Female guts were likely to have higher levels of *Staphylococcus and Pseudomonas*, whereas males had higher levels of *Enterococcus* (**Figure 2D**). Moreover, their relative importance vis-à-vis infection and immunity varied across the sexes. For instance, disrupting the microbiota with antibiotics improved post-infection survival in females (from 50% to ∼75%; **Figure 2E, Table S6**) but had no significant effect in males, thereby eliminating the observed difference in infection susceptibility between the sexes (**Fig 2E**). Antibiotic-fed females also showed reduced pathogen load (**Figure 2F, Table S5**), indicating improved pathogen resistance. Reintroducing microbiota to antibiotic-fed beetles rescued the phenotypic divergence in both infection susceptibility and bacterial load (**Figure 2G, H; Table S6, 5**). Finally, six days of initial antibiotic exposure played a crucial role in the loss of sexual dimorphism because transferring antibiotic-treated beetles subsequently to faeces-free autoclaved wheat bran for the following six days (i.e. treatment D) did not rescue the female survival (**Figure S2, Table S7**).

We also confirmed that antibiotic treatment successfully depleted the gut microbiota in both sexes as the gut bacterial DNA isolated from the beetles was below the detection level of PCR amplification (**Figure S3**). We were successful in reconstituting the autoclaved wheat bran by pre-conditioning it with adults and larvae as evident from broad patterns of similarity seen between microbiota-reconstituted wheat bran and normal wheat bran (**Figure 2D**). Diet-mediated reconstitution of the microbiome of antibiotic-fed females was also highly effective in reconstituting the gut bacteria. However, the composition appeared to be different from that of standard beetles used as procedural control— e.g., unlike females raised in the standard diet, females feeding on conditioned wheat bran showed a higher abundance of *Bacillus* and *Virgibacillus sp*, whereas the abundance of bacterial genera *Staphylococcus* and *Pseudomonas* was low (**Figure 2D**). We could not detect microbiota in reconstituted males, possibly indicating sexual divergence in the rate of feeding or acquisition of microbes within the experimental window. Individual replicates within the same treatment showed qualitatively similar patterns of microbiota composition (see **Figure S4**).

### Individual immune components do not consistently explain the observed phenotypic variations

Finally, we investigated a few key components of beetle immunity to explain their sexually divergent infection susceptibility and bacterial load. Although control females reared under a standard diet showed a lower expression of tenecin 1, compared to males (**Figure 3A, Table S8**), such sex-specific gene expression pattern was not found in beetles fed on antibiotic-supplemented or microbiota-reconstituted beetles (**Figure 3B, C, Table S8)**. We also failed to find any sex differences in the expression levels of AMP tenecin *4* (**Figure S5, Table S9**) or PO activity (**Figure S6, Table S10**), regardless of their microbiota-manipulation status.

**Figure 3:**
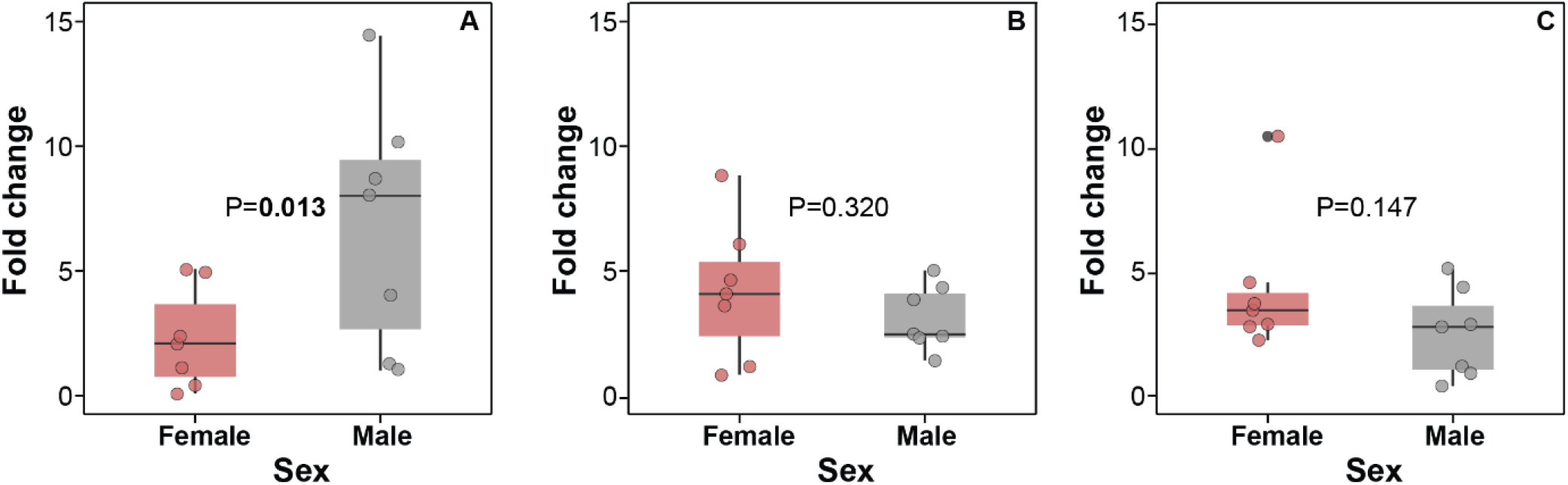
(**A–C**) Relative fold-change in tenecin 1 expression at 8 hours post-infection with 0.25 OD Bt (720 ± 50 cells/beetle) across sexes and different dietary treatments (**A**-fed on a normal diet; **B**-fed on an antibiotic-supplemented diet; **C**-fed on a microbiota-reconstituted diet) (n=7/sex/dietary treatment). In each panel, P-values represent the effect of sex on gene expression (data analysed using a generalised linear model fitted to a Gamma distribution).

## DISCUSSION

Our experiments revealed critical insights into the role of microbiota in maintaining differences in immunity and infection susceptibility between male and female *T. molitor* beetles. Typically, females were more susceptible to bacterial infections and had a higher bacterial load, but these differences disappeared when their gut microbiota was depleted with antibiotics. Intriguingly, reintroducing the microbiome through feeding on conditioned wheat bran could rescue the sexually divergent survival patterns and bacterial load variations, indicating the causative role of the microbiome on sex-dependent infection outcomes. Notably, only female beetles responded to microbiota changes, whereas males remained unaffected. The importance of microbiota in infection responses thus varied between sexes, presenting an exciting area for further exploration of the underlying mechanistic links.

Our finding of improved survival of experimental females after microbiome depletion aligns with previous studies where microbiota had a causative role in increasing disease susceptibility (26, 27). For example, eliminating gut microbiota in different lepidopteran species, including *Manduca sexta*, could enhance resistance to infections with *B. thuringiensis* and its toxins (26). Conversely, higher microbiome diversity or probiotic intake may exacerbate vulnerability to parasites in bumble bees (28) or increase the parasitic load in mice (29). The elevated risk of pathogen susceptibility in these studies is thus a possible trade-off for engaging with microbiota. However, it is presently unclear why such cost was restricted to only females in our experiments.

Additionally, our study hinted at notable divergence in gut microbiota profiles between females reared under a normal vs reconstituted (with faecal matter) wheat bran despite having similar infection susceptibility and pathogen load. This difference might stem from the fact that the reconstituted wheat bran microbiota consisted of a blend of microbial communities from both larval and adult beetles, possibly creating a distinct combination of diet microbiota compared to those naturally present in the adult gut (30). Nonetheless, this indeed prompts questions about the relative importance of overall microbiome diversity versus the presence of specific taxa in modulating infection susceptibility and compromising host health in our experimental females. Interestingly, both groups of females shared the *Enterococcus* species, which is already recognised for providing resistance to *B. thuringiensis* in a related flour beetle species (31). Additionally, highly prevalent *Bacillus* species in reconstituted female guts (showing an overall abundance of ∼70%) may also aid in reviving the original survival patterns potentially via oral priming effects (32), but more investigation is necessary to confirm these causal links. Also, we could only assay bacterial communities, whereas the role of fungal or viral communities remains unexplored.

Microbiota also played a key role in determining the association between immune gene expression and infection outcomes in our experiments. For example, females had reduced tenecin-1 levels than males, which could explain why they could not control the pathogen growth as effectively as males and, hence, were more vulnerable to *B. thuringiensis* infection under normal conditions. However, both antibiotic-supplemented or microbiota-reconstituted females did not show changes in tenecin-1 levels despite having contrasting post-infection survival and bacterial load variations. Conversely, it was surprising that males reduced their tenecin-1 levels after microbiota depletion, which led to the loss of sexually dimorphic gene expression, but without any effects on their post-infection survival. These results with males mirrored previous studies where animals reared under axenic conditions reduced the expression of several immune-related transcripts (33, 34). However, more studies are needed to pinpoint why the functional association between microbiota and immune genes varied across sexes. Also, we observed no changes in phenoloxidase activity (13) or the expression of another AMP called tenecin-4, which is typically reactive to Gram-negative bacteria (24). Thus, alterations in the beetles’ microbiota in our experiments may have invoked different immune mechanisms unrelated to AMPs and melanisation while combating pathogens.

In conclusion, our research highlights the critical role of the microbiome in regulating sexually dimorphic immunity, infection susceptibility, and pathogen load in an insect model. However, joint analyses of different components of the insect immune system are perhaps necessary to understand how their sex-specific associations with microbiota modulate sexually dimorphic pathogen growth dynamics and infection susceptibility. Finally, it is vital to parallelly investigate the link between the microbiome and various host life-history traits, including effects on development and reproduction, to reveal how gut bacteria, which are widely considered beneficial to organismal fitness (35), can turn harmful in the presence of pathogenic infections. This will provide a broader understanding of host-microbiota coevolutionary adaptations impacting health and disease vulnerability.

## AUTHOR’S CONTRIBUTIONS

Conceptualisation: Imroze Khan, Srijan Seal

Design of the experiment: Imroze Khan, Srijan Seal, Pavankumar Thunga, Rhitoban Raychoudhury

Data curation: Srijan Seal, Manisha Gupta,

Formal analysis: Srijan Seal, Manisha Gupta

Funding acquisition: Imroze Khan, Rhitoban Raychoudhury

Investigation: Srijan Seal, Devashish Kumar, Pavankumar Thunga, Pawan Khangar, Manisha Gupta, Dipendra Nath Basu

Supervision: Imroze Khan

Visualisation: Srijan Seal, Manisha Gupta

Writing – original draft: Srijan Seal, Imroze Khan

Writing – review & editing: Imroze Khan, Srijan Seal,

## ACKNOWLEDGEMENTS

We acknowledge Biswajit Shit for his feedback on the manuscript.

## REFERENCES

1. C. D. Kelly, A. M. Stoehr, C. Nunn, K. N. Smyth, Z. M. Prokop, Sexual dimorphism in immunity across animals: a meta-analysis. Ecol. Lett. 21, 1885–1894 (2018).

2. S. L. Klein, K. L. Flanagan, Sex differences in immune responses. Nat. Rev. Immunol. 16, 626– 638 (2016).

3. C. L. Nunn, P. Lindenfors, E. R. Pursall, J. Rolff, On sexual dimorphism in immune function. Philos. Trans. R. Soc. B Biol. Sci. 364, 61–69 (2009).

4. M. Zuk, The sicker sex. PLoS Pathog. 5, e1000267 (2009).

5. R. L. Belmonte, M.-K. Corbally, D. F. Duneau, J. C. Regan, Sexual dimorphisms in innate immunity and responses to infection in Drosophila melanogaster. Front. Immunol. 10, 3075 (2020).

6. D. Duneau, P. Luijckx, L. F. Ruder, D. Ebert, Sex-specific effects of a parasite evolving in a female-biased host population. BMC Biol. 10, 104 (2012).

7. C. D. Kelly, B. R. Tawes, Sex-specific effect of juvenile diet on adult disease resistance in a field cricket. PLoS ONE 8, e61301 (2013).

8. O. Cervantes, et al., Role of hormones in the pregnancy and sex-specific outcomes to infections with respiratory viruses. Immunol. Rev. 308, 123–148 (2022).

9. R. Vemuri, et al., The microgenderome revealed: sex differences in bidirectional interactions between the microbiota, hormones, immunity and disease susceptibility. Semin. Immunopathol. 41, 265–275 (2019).

10. M. Elderman, et al., Sex and strain dependent differences in mucosal immunology and microbiota composition in mice. Biol. Sex Differ. 9, 26 (2018).

11. F. Fransen, et al., The impact of gut microbiota on gender-specific differences in immunity. Front. Immunol. 8, 754 (2017).

12. Junrui-Fu, et al., Gut dysbacteriosis induces expression differences in the adult head transcriptome of Spodoptera frugiperda in a sex-specific manner. BMC Microbiol. 23, 388 (2023).

13. I. Khan, D. Agashe, J. Rolff, Early-life inflammation, immune response and ageing. Proc. R. Soc. B Biol. Sci. 284, 20170125 (2017).

14. P. Bolme, M. Eriksson, D. Habte, L. Paalzow, Pharmacokinetics of streptomycin in Ethiopian children with tuberculosis and of different nutritional status. Eur. J. Clin. Pharmacol. 33, 647– 649 (1988).

15. Y. Yokoyama, et al., The pharmacokinetics of ampicillin–sulbactam in anuric patients: dosing optimization for prophylaxis during cardiovascular surgery. Int. J. Clin. Pharm. 38, 771–775 (2016).

16. C. J. Mason, et al., Diet influences proliferation and stability of gut bacterial populations in herbivorous lepidopteran larvae. PLOS ONE 15, e0229848 (2020).

17. R. Agarwal, M. Gupta, A. Antony, R. Sen, R. Raychoudhury, In vitro studies reveal that Pseudomonas, from Odontotermes obesus colonies, can function as a defensive mutualist as it prevents the weedy fungus while keeping the crop fungus unaffected. Microb. Ecol. 84, 391– 403 (2022).

18. M. C. Frith, M. Hamada, P. Horton, Parameters for accurate genome alignment. BMC Bioinformatics 11, 80 (2010).

19. I. Khan, A. Prakash, D. Agashe, Divergent immune priming responses across flour beetle life stages and populations. Ecol. Evol. 6, 7847–7855 (2016).

20. M. Mulet, M. Gomila, B. Lemaitre, J. Lalucat, E. García-Valdés, Taxonomic characterisation of Pseudomonas strain L48 and formal proposal of Pseudomonas entomophila sp. nov. Syst. Appl. Microbiol. 35, 145–149 (2012).

21. T. Therneau, A package for survival analysis in R. (2024). Deposited 13 February 2024.

22. H. Tabunoki, N. T. Dittmer, M. J. Gorman, M. R. Kanost, Development of a new method for collecting hemolymph and measuring phenoloxidase activity in Tribolium castaneum. BMC Res. Notes 12, 7 (2019).

23. C. Zanchi, P. R. Johnston, J. Rolff, Evolution of defence cocktails: Antimicrobial peptide combinations reduce mortality and persistent infection. Mol. Ecol. 26, 5334–5343 (2017).

24. J.-H. Chae, et al., Purification and characterization of tenecin 4, a new anti-Gram-negative bacterial peptide, from the beetle Tenebrio molitor. Dev. Comp. Immunol. 36, 540–546 (2012).

25. T. D. Schmittgen, K. J. Livak, Analyzing real-time PCR data by the comparative CT method. Nat. Protoc. 3, 1101–1108 (2008).

26. N. A. Broderick, et al., Contributions of gut bacteria to Bacillus thuringiensis-induced mortality vary across a range of Lepidoptera. BMC Biol. 7, 11 (2009).

27. R. Visweshwar, H. C. Sharma, S. M. D. Akbar, K. Sreeramulu, Elimination of gut microbes with antibiotics confers resistance to Bacillus thuringiensis toxin proteins in Helicoverpa armigera (Hubner). Appl. Biochem. Biotechnol. 177, 1621–1637 (2015).

28. D. P. Cariveau, J. Elijah Powell, H. Koch, R. Winfree, N. A. Moran, Variation in gut microbial communities and its association with pathogen infection in wild bumble bees (Bombus). ISME J. 8, 2369–2379 (2014).

29. M. A. Dea-Ayuela, S. Rama-Iñiguez, F. Bolás-Fernandez, Enhanced susceptibility to Trichuris muris infection of B10Br mice treated with the probiotic Lactobacillus casei. Int. Immunopharmacol. 8, 28–35 (2008).

30. C. Manthey, P. R. Johnston, S. Nakagawa, J. Rolff, Complete metamorphosis and microbiota turnover in insects. Mol. Ecol. 32, 6543–6551 (2023).

31. T. Grau, A. Vilcinskas, G. Joop, Probiotic Enterococcus mundtii Isolate protects the model insect Tribolium castaneum against Bacillus thuringiensis. Front. Microbiol. 8, 1261 (2017).

32. J. M. Greenwood, et al., Oral immune priming with Bacillus thuringiensis induces a shift in the gene expression of Tribolium castaneum larvae. BMC Genomics 18, 329 (2017).

33. L. Auger, S. Bouslama, M.-H. Deschamps, G. Vandenberg, N. Derome, Absence of microbiome triggers extensive changes in the transcriptional profile of Hermetia illucens during larval ontogeny. Sci. Rep. 13, 2396 (2023).

34. K. J. Vogel, L. Valzania, K. L. Coon, M. R. Brown, M. R. Strand, Transcriptome sequencing reveals large-scale changes in axenic Aedes aegypti larvae. PLoS Negl. Trop. Dis. 11, e0005273 (2017).

35. D. N. Lesperance, N. A. Broderick, Microbiomes as modulators of Drosophila melanogaster homeostasis and disease. Curr. Opin. Insect Sci. 39, 84–90 (2020).

