## Supplementary file for "MICROBIOTA DRIVES THE SEXUALLY DIMORPHIC INFECTION OUTCOMES IN MEALWORM BEETLES"

**SUPPLEMENTARY METHODS**

**Quantifying the changes in the microbiome composition**

We characterized the microbiome composition of beetles maintained under different dietary treatments using pool sequencing as well as individual beetle separately to get a comprehensive understanding of microbial diversity. We amplified a portion of 16S rRNA gene from the isolated microbial DNA from the gut of beetle samples using universal primers (1) (Table S1). We had three biological replicates for each of the treatments (Males and females fed on normal wheat bran (Treatment A) and females fed on reconstituted diet (Treatment C). We setup three separate PCR reactions (technical replicates) for each biological replicate. Finally, we pooled equimolar concentrations of the three PCR reactions of three biological replicates into one tube for downstream processing. We used the SQK-LSK 108 ligation sequencing kit (Oxford Nanopore technologies) for Nanopore library preparation (2). We polyadenylated each sample using the Ultra II end preparation reaction mix (Oxford Nanopore Technologies). We then cleaned these samples using the AMPure XP beads and eluted in sterile nuclease-free water. Next, we ligated barcode adaptors for the addition of unique barcodes using the EXP-PBC001 PCR barcoding kit (Oxford Nanopore Technologies). We purified these barcoded PCR products using the QIAquick PCR purification kit and quantified using a Qubit 3.0 Fluorometer (Life Technologies). We pooled all the barcoded samples in equimolar concentrations to prepare a final pooled library amounting to 1μg. Finally, this pooled library was further polyadenylated for the addition of sequencing adaptors and cleaned using the AMPure XP beads. We also added 5μl of λ phage DNA as a control. We sequenced the samples on the MinION Flow cell (FLO-MIN106) R9 version with the protocol “NC_48Hr_sequencing_FLO-MIN106_SQK-LSK108_plus_Basecaller” for 48hrs. After sequencing, we converted raw reads from FAST5 to FASTQ (‘basecalled’) and separated according to barcodes (demultiplexed) using the Guppy program (Oxford Nanopore Technologies Ltd). Next, we subjected FASTQ reads to the Nanofilt program (Oxford Nanopore Technologies Ltd) to filter high-quality reads having an average score ≥10. Next, we trimmed sequencing reads as per target read length for 16S primers. We only processed high-quality trimmed reads for taxonomic identification. To determine bacterial community composition in each sample, we identified the reads using the Lastal program in LAST v973 (3) by aligning these queries against the NCBI reference database with the following parameters: gap opening penalty of 1, match score of 1, and gap extension penalty of 1 (4).

For sequencing the individual biological samples in each treatment separately, we sent the isolated DNA to an external sequencing facility (miBiome therapeutics). The isolated DNA samples was amplified using universal primers as mentioned previously. The PCR products were purified followed by adaptor ligation to generate libraries following Illumina Inc. protocol. The cleaned libraries were quantified on Qubit fluorometer and appropriate dilutions were loaded on HS D1000 screen tape to determine the size range of fragments and the average library size. The samples were sequenced on Illumina Miseq platform. We analyzed the qualities of the sequenced raw reads using fastqc (v0.12.1). We use cutadapt (v 4.8) for removing primer sequences and for filtering. Subsequently we implemented a standalone algorithm DADA2 (Divisive Amplicon Denoising Algorithm 2) for further trimming and truncating using the recommended parameters (5). We generated an error model for both the forward and reverse reads for denoising measures. We merged paired reads and performed a sanity check for chimeric sequences. We used databases like SILVA (silva_nr99_v138.1), RDP (rdp_train_set_18) and GTDB (GTDB_bac120_arc53_ssu_r207) (6) to assign taxonomy using the assigntaxonomy function in DADA2. We further validated the taxonomic classification by using BLASTn search against the nucleotide database of NCBI. We created a phyloseq object in R using the count and taxonomy files with the help of the phyloseq package (v 1.44.0). We excluded the reads which were classified as mitochondria and chloroplast from the analysis and calculated the relative abundance of all the taxa in R version 4.3.1.

**B. Assaying post-infection survival**

We initiated overnight cultures of *Bacillus thuringiensis* (Bt) or *Pseudomonas entomophila* (Pe) from glycerol stocks by growing in liquid LB media at 30°C in shaking incubators. We prepared the infection dose by diluting a secondary culture of Bt (or Pe) that was allowed to reach 1 OD before centrifuging at 4000 rpm at 4 degrees for 5 minutes and washing the obtained pellet three times with sterile Ringer solution (7). We used three infection doses for Bt which were 0.25 OD (720 ± 50 cells/beetle), 0.5 OD (1320 ± 140 cells/beetle) and 1 OD (2125 ± 180 cells/beetle. For Pe infection, we used a single dose of 0.05 OD (6700 ± 850 cells/beetle)). We infected beetles with different doses of the Gram-positive bacteria *Bacillus thuringiensis* DSM 2046 (Bt) or with a single dose of *Pseudomonas entomophila* (Pe) by anaesthetizing them on ice for 5 minutes followed by injecting 5 µl of Bt (or Pe) cells directly between the 3rd and 4th abdominal sternite using a fine glass capillary tube. We injected 5 µl of sterile Ringer solution into a subset of beetles as procedural control. We returned the beetles to individual chambers of 12-well plates post-infection (or mock infection) with ad libitum food (8) and monitored daily for any mortality. We used Cox proportional hazard model for analysing post-infection survival.

**C. Measuring the Bt load across the infection window**

We measured the load of Bt cells in the haemolymph of infected beetles at three time points post-infection (5 hrs, 10 hrs and 20 hrs) following a previously published protocol (8). Briefly, we made an incision on the genitalia of anaesthetized beetles with a fine scalpel and placed each beetle in a 0.5 ml microcentrifuge tube with a pinhole made at the bottom. The microcentrifuge tube was then placed in a 1.5 ml microcentrifuge tube containing 100µl of 1X PBS and centrifuged at 6,000 rpm for 10 minutes at 4°C.  The haemolymph was collected in the 1.5ml microcentrifuge tube. 400µl of 1X PBS was added to the tube and diluted serially to 10^-2^ fold. 20µl from each serial dilution was spread plated on Luria Bertani agar using glass beads. We counted colonies from the first dilution that had isolated CFU’s after overnight incubation. We analyzed the bacterial load data using a generalized linear model fitted to a gamma distribution with sex and time as fixed effects.

**D.** **Measuring changes in fold expression of immune transcripts**

We dissected the fat body from abdominal section of the beetles using fine scalpels and tweezers (Fine Science tools). We isolated RNA using Trizol reagent (Ambion), following the manufacturer’s protocol. The isolated RNA was further purified using Turbo DNA-free kit (Ambion) to digest any remaining DNA. cDNA was synthesized using iScript cDNA synthesis kit (Bio-Rad), following the manufacturer’s protocol. We performed qPCR in a step one real-time thermocycler (QuantStudio 5 Applied Biosystems) using SYBR green (from kappa) with 500 ng of cDNA as template per 10µl reaction volume (9). We calculated the ΔC_t_ for gene of interest for individual sample by subtracting the mean C_t_ value of target gene (tenecin 1 and tenecin 4) from mean C_t_ value of reference gene rpl27a (a ribosomal protein subunit previously shown to be stable after infection) (10). We calculated the relative change in fold expression level as 2^-ΔΔCt^ where ΔΔC_t_ is the difference in ΔC_t_ values between infected individual and sham infected individual (11). We measured the relative fold change in gene expression using a generalized linear model fitted to Gamma distribution.

**E. Assaying the phenoloxidase activity**

We measured the phenoloxidase activity following a previously published protocol (6). We extracted 5µl of haemolymph by inducing a wound between the head and the thorax from infected (or mock-infected) beetles two hours post-infection and added it to a microcentrifuge tube containing 20µl of 1X PBS. We centrifuged the sample at 6000 rpm for 10 minutes at 4 °C to remove cell debris. We transferred 5 µl of the supernatant to a 96-well-microplate kept on ice containing 140µl distilled water and 20µl of 1X PBS. Next, we added 20µl of L-DOPA substrate into each well and immediately placed the plate in a microplate reader. We tracked the rate of formation of dopachrome by measuring the absorbance at 490 nm every minute for a period of 1 hr at 30°C. We measured PO enzyme activity (V_max_) as the slope of the linear phase of the reaction. We analyzed the differences between the slopes using Wilcoxon rank sum test.

**References**

1.	W. Zheng, *et al.*, An accurate and efficient experimental approach for characterization of the complex oral microbiota. *Microbiome* **3**, 48 (2015).

2.	R. Agarwal, M. Gupta, A. Antony, R. Sen, R. Raychoudhury, In vitro studies reveal that *Pseudomonas*, from *Odontotermes obesus* colonies, can function as a defensive mutualist as it prevents the weedy fungus while keeping the crop fungus unaffected. *Microb. Ecol.* **84**, 391–403 (2022).

3.	M. C. Frith, M. Hamada, P. Horton, Parameters for accurate genome alignment. *BMC Bioinformatics* **11**, 80 (2010).

4.	J. Shin, *et al.*, Analysis of the mouse gut microbiome using full-length 16S rRNA amplicon sequencing. *Sci. Rep.* **6**, 29681 (2016).

5.	L. Kešnerová, *et al.*, Gut microbiota structure differs between honeybees in winter and summer. *ISME J.* **14**, 801–814 (2020).

6.	A. R. Odom, T. Faits, E. Castro-Nallar, K. A. Crandall, W. E. Johnson, Metagenomic profiling pipelines improve taxonomic classification for 16S amplicon sequencing data. *Sci. Rep.* **13**, 13957 (2023).

7.	I. Khan, D. Agashe, J. Rolff, Early-life inflammation, immune response and ageing. *Proc. R. Soc. B Biol. Sci.* **284**, 20170125 (2017).

8.	H. Tabunoki, N. T. Dittmer, M. J. Gorman, M. R. Kanost, Development of a new method for collecting hemolymph and measuring phenoloxidase activity in *Tribolium castaneum*. *BMC Res. Notes* **12**, 7 (2019).

9.	Y. H. Jo, *et al.*, TmCactin plays an important role in Gram-negative and -positive bacterial infection by regulating expression of 7 AMP genes in *Tenebrio molitor*. *Sci. Rep.* **7**, 46459 (2017).

10.	C. Zanchi, P. R. Johnston, J. Rolff, Evolution of defence cocktails: Antimicrobial peptide combinations reduce mortality and persistent infection. *Mol. Ecol.* **26**, 5334–5343 (2017).

11.	T. D. Schmittgen, K. J. Livak, Analyzing real-time PCR data by the comparative CT method. *Nat. Protoc.* **3**, 1101–1108 (2008).

**SUPPLEMENTARY FIGURES**

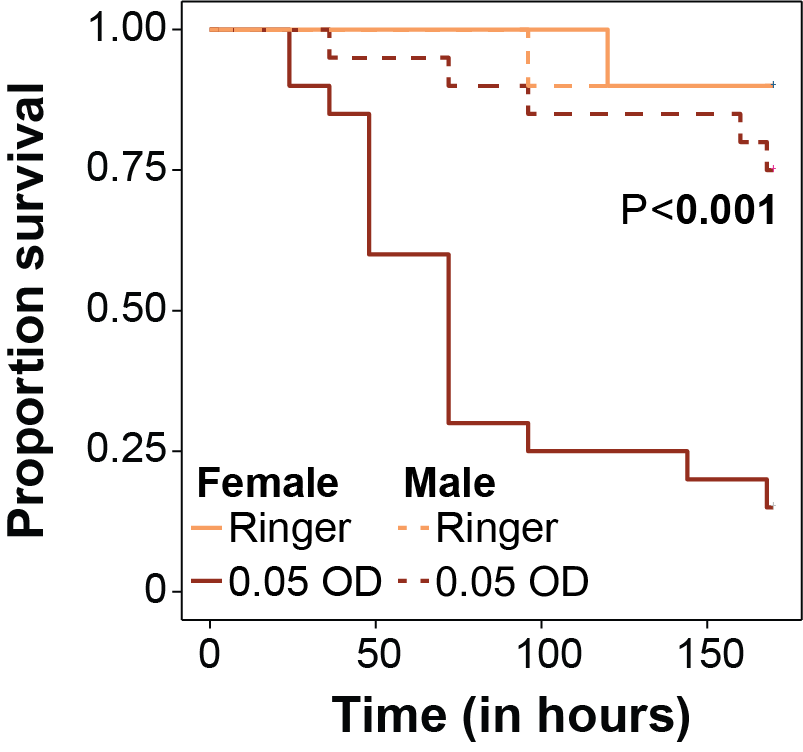

**Figure S1:** Proportion of beetles surviving after infection with *Pseudomonas entomophila* (Pe) (n=20 beetles/sex) under normal dietary conditions. The P value represents the effect of sex on post-infection survival (data analysed using the Cox proportional hazard model).

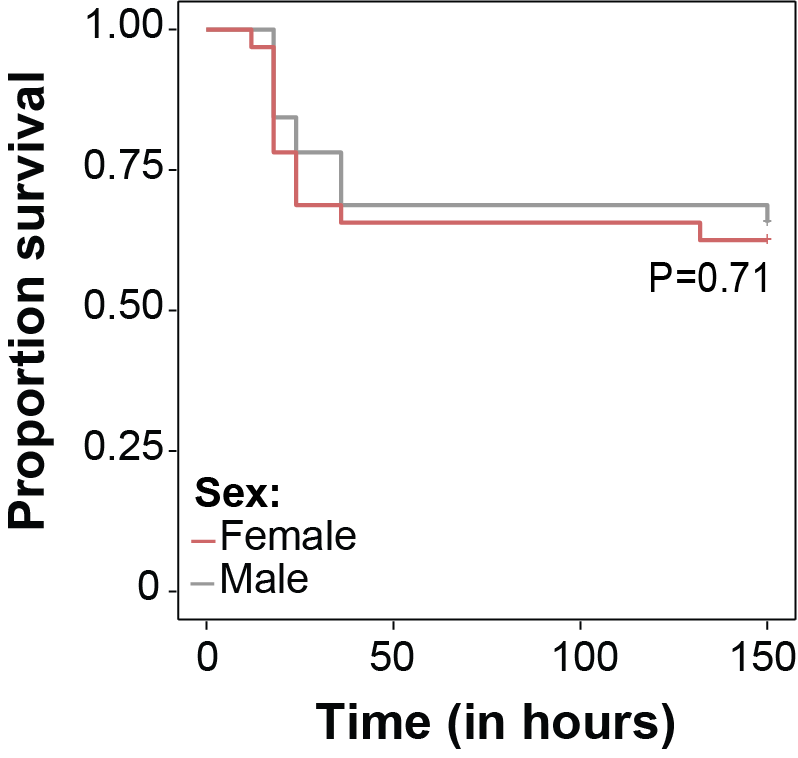

**Figure S2:** Post-infection survival of beetles exposed to an antibiotic-treated diet for six days followed by another six days exposure to faeces-free autoclaved wheat bran (n=32 beetles/sex) (i.e., shorter exposure to antibiotics; see Treatment D in main text). The P value represents the effect of sex on post-infection survival (data analysed using the Cox proportional hazard model).

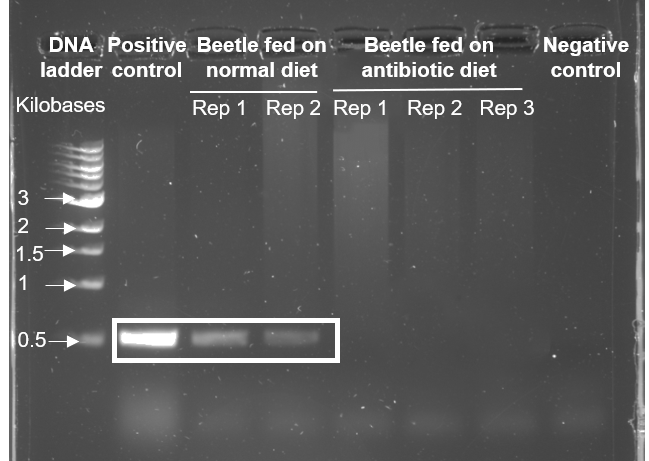

**Figure S3:** A representative image of 1% agarose gel stained with SYBR safe showing the amplified PCR products of V3-V4 region of 16S rRNA gene of microbes. Lane 1 contains the 1kb DNA ladder (New England Biolabs). Lane 2 contains the amplified product obtained from the isolated *Escherichia coli* genomic DNA (positive control). Lanes 3–4 are amplified products obtained from beetles fed on a normal diet (2 replicates). Lanes 5–7 contain samples obtained from beetles fed on the antibiotic-supplemented diet (3 replicates). Lane 8 is for the negative control. The amplified products (~500bp) are highlighted.

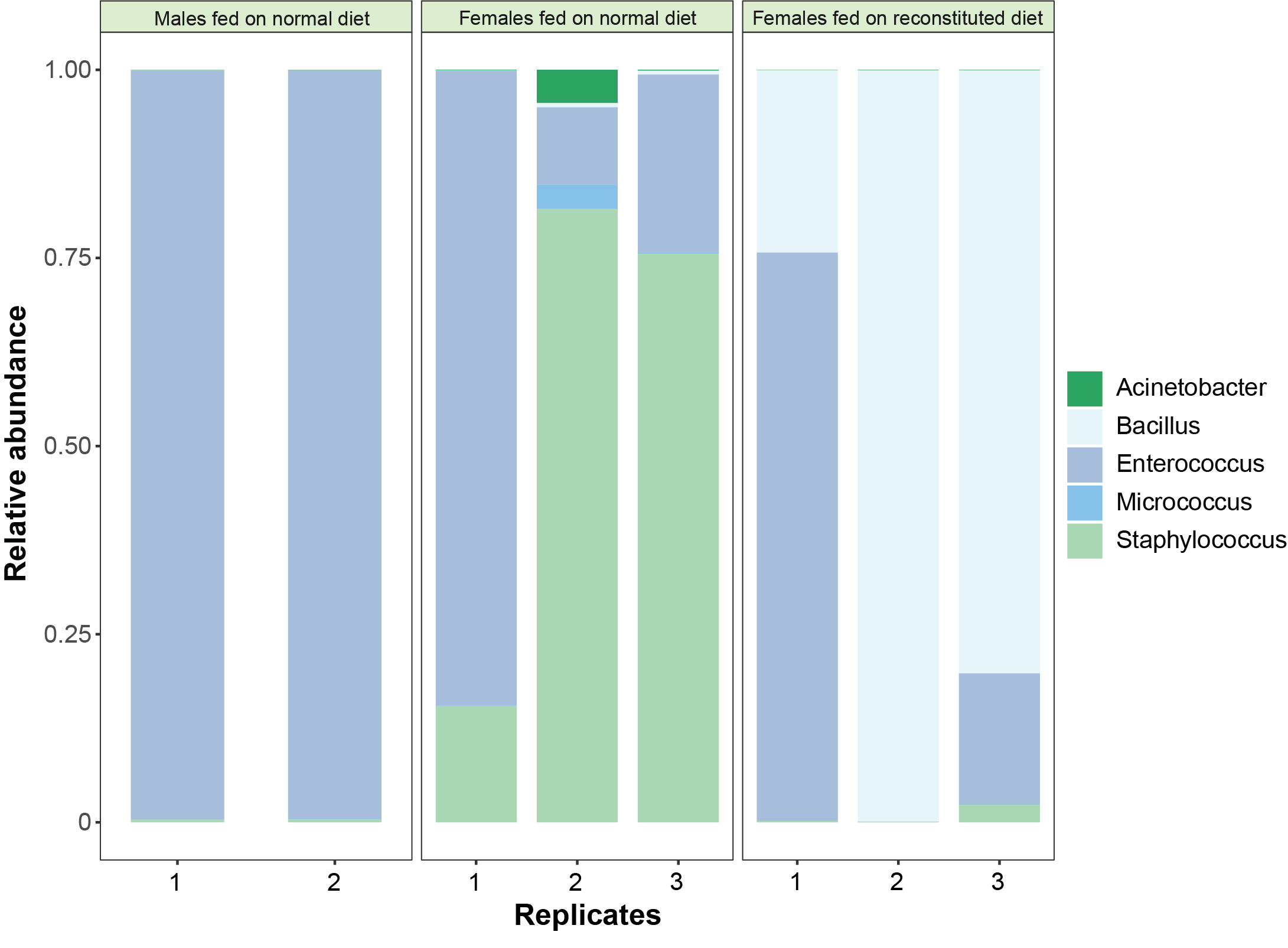

**Figure S4:** Stacked bar plots showing the relative abundance of the five most abundant bacterial amplicon sequence variants separately in each replicate beetle within each treatment (N=2 for males fed on a normal diet and N=3 for females fed on normal diet or microbiota-reconstituted diet). We lost one replicate of males fed on a normal diet due to experimental error.

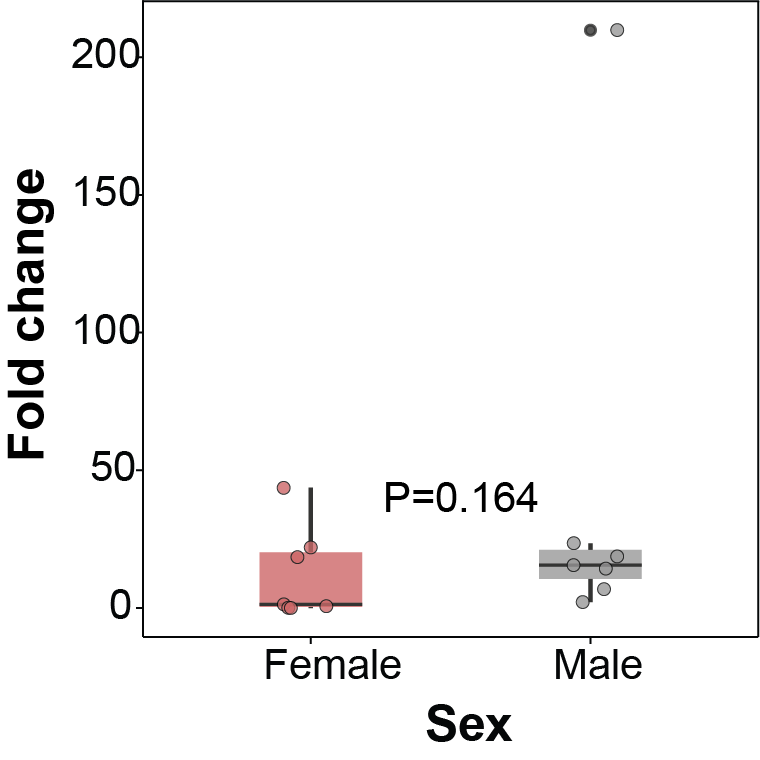

**Figure S5:** Relative fold-change in *tenecin* 4 expression at 8 hours post-infection with 0.25 OD Bt across sexes when fed on a normal diet (n=7 beetles/sex). The P value represents the effect of sex on fold-changes of gene expression (data analysed using a generalised linear model fitted to a Gamma distribution).

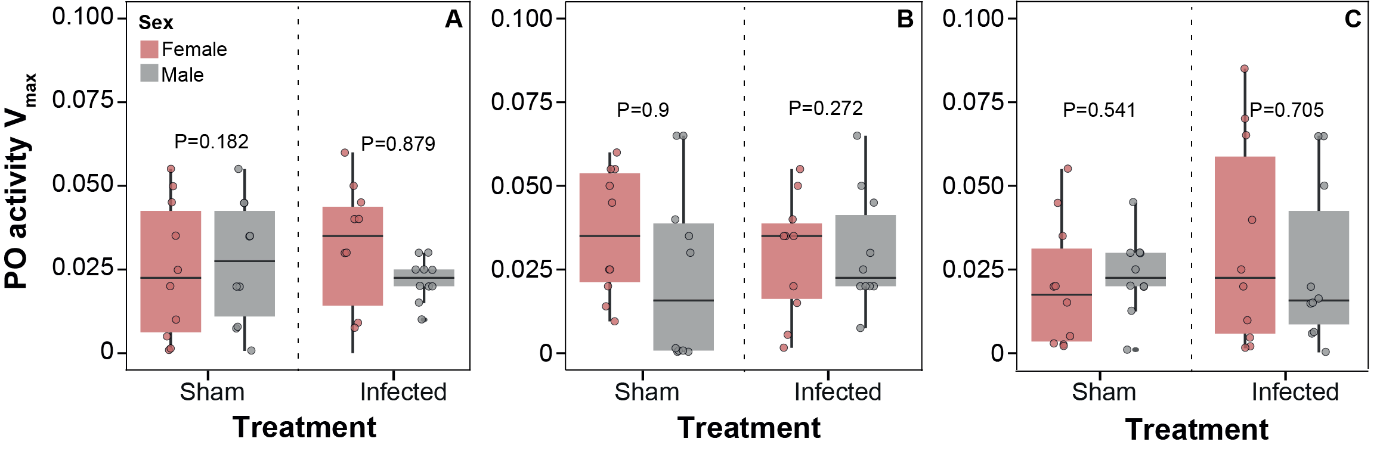

**Figure S6:** Phenoloxidase activity (PO) after Bt infection in beetles fed on a (A) normal diet, (B) antibiotic-supplemented diet and (C) microbiota-reconstituted diet (C)**.** PO activity was measured as V_max_ of the enzymatic reaction (slope of the linear phase of the reaction curve) (n= 10 beetles/infection treatment/sex). In each panel, P values represent the effect of sex within each infection treatment across dietary treatments (data analysed using the Wilcoxon rank sum test).

**Table S1. Sequences of primers used for amplifying 16S rRNA V3-V4 region of isolated microbial DNA obtained from Zheng et al.** (1)**.**

| **Name of primer** | **Sequence** |
| --- | --- |
| Illumina_16S_341F | TCGTCGGCAGCGTCAGATGTGTATAAGAGACAGCCTACGGGNGGCWGCAG |
| Illumina_16S_806R | GTCTCGTGGGCTCGGAGATGTGTATAAGAGACAGGACTACHVGGGTATCTAATCC |

**Table S2. Sequences of qPCR primers used for measuring expression levels of antimicrobial peptides and house-keeping gene obtained from Zanchi et al.** (10)**.**

| **Gene** | **Forward primer 5’-3’** | **Reverse primer 5’-3’** |
| --- | --- | --- |
| *Tenecin 1* | GGAAGCGGCAACAGCTGAAGAAAT | AACGCAGACCCTCTTTCCGTTACA |
| *Tenecin 4* | TCA ACAACGGCGGCCACAAATTAGA | TCTTCGGTGGGAAGCTGGATTACA |
| *Rpl27a* | TCGGAAAGTTGGGAATGAGG | TTTGACCTTGTCTGCTCACT |

**Table S3.** Summary of the Cox proportional hazard analysis of survival data after infection with different doses of *Bacillus thuringiensis* (Bt), using sex (S) and infection dose (D) as fixed effects. [Model specification: Survival ~ Sex + Infection dose + Sex $\times$ Infection dose]. Significant effects are highlighted in bold.

| **Infection** | **Effects** | **Loglikelihood** | **χ^2^** | **df** | **P** |
| --- | --- | --- | --- | --- | --- |
|  | S | -326.68 | 12.285 | **1** | **<0.001** |
| Bt | D | -304.44 | 44.485 | 3 | **<0.001** |
|  | S$\times$ D | -301.48 | 5.905 | 3 | 0.116 |

**Table S4.** Summary of the Cox proportional hazard analysis of survival data after infection with *Pseudomonas entomophila* (Pe) with sex as a fixed effect. [Model specification: Survival ~ Sex]. Significant effects are highlighted in bold.

| **Infection** | **Effects** | **Loglikelihood** | **χ^2^** | **df** | **P** |
| --- | --- | --- | --- | --- | --- |
| Pe | Sex | -65.38 | 17.07 | **1** | **<0.001** |

**Table S5.** Summary of a generalised linear model on log-transformed bacterial load data, fitted to a Gamma distribution, as a function of sex (S) and time-points of the assay (T) [Model specification: Bacterial load ~ Sex + Time point + Sex $\times$ Time point]. Significant effects are highlighted in bold.

| **Treatment** | **Effects** | **χ^2^** | **df** | **P** |
| --- | --- | --- | --- | --- |
| Treatment A (fed on a normal diet) | S | 3.0059 | 1 | 0.083 |
|  | T | 1.7224 | 2 | 0.423 |
|  | S $\times$ T | 1.1771 | 2 | 0.555 |
| Treatment B (fed on an antibiotic- supplemented diet) | S | 1.0718 | 1 | 0.301 |
|  | T | 1.3151 | 2 | 0.518 |
|  | S $\times$ T | 3.9731 | 2 | 0.137 |
| Treatment C (fed on a microbiota-reconstituted diet) | S | 7.3122 | 1 | **0.007** |
|  | T | 7.1230 | 2 | **0.029** |
|  | S $\times$ T | 1.1405 | 2 | 0.565 |

**Table S6.** Summary of a mixed-effects Cox model on survival data after infection with *Bacillus thuringiensis* (Bt) within each dietary treatment (Treatment A, B, C) to manipulate the microbiome, with sex as a fixed effect and replicated trials as a random effect. [Model specification: Survival ~ Sex +(1|trial)]. Significant effects are highlighted in bold.

| **Treatment** | **Effect** | **Loglikelihood** | **χ^2^** | **df** | **P** |
| --- | --- | --- | --- | --- | --- |
| Treatment A (fed on a normal diet) | Sex | -550.13 | 33.092 | **1** | **<0.001** |
| Treatment B (fed on an antibiotic-supplemented diet) | Sex | -248.46 | 3.8265 | 1 | 0.054 |
| Treatment C (fed on a microbiota-reconstituted diet) | Sex | -575.72 | 30.65 | 1 | **<0.001** |

**Table S7**. Response to Bt infection after feeding on antibiotic-supplemented wheat bran followed by autoclaved wheat bran (Treatment C). The table provides a summary of Cox proportional hazard analyses of survival data after infection with *Bacillus thuringiensis* (Bt) with sex as fixed effects. [Model specification: Survival ~ Sex].

| **Infection** | **Effects** | **Loglikelihood** | **χ^2^** | **df** | **P** |
| --- | --- | --- | --- | --- | --- |
| Bt | Sex | -91.064 | 0.139 | **1** | 0.709 |

**Table S8.** Summary of generalised linear models on post-infection fold changes of *tenecin* 1 expression level, fitted to a Gamma distribution, as a function of sex within each dietary treatment [Model specification: Fold change~ Sex]. Significant effects are highlighted in bold.

| **Treatment** | **Effect** | **χ^2^** | **df** | **P** |
| --- | --- | --- | --- | --- |
| Treatment A (fed on a normal diet) | Sex | 6.0663 | 1 | **0.013** |
| Treatment B (fed on an antibiotic-supplemented diet) | Sex | 0.9885 | 1 | 0.320 |
| Treatment C (fed on a microbiota-reconstituted diet) | Sex | 2.1037 | 1 | 0.147 |

**Table S9.** Summary of a generalised linear model on post-infection fold changes of *tenecin* 4 expression level, fitted to a Gamma distribution, as a function of sex in beetles fed on normal diet [Model specification: Fold change~ Sex].

| **Effect** | **χ^2^** | **df** | **P** |
| --- | --- | --- | --- |
| Sex | 1.933 | 1 | 0.164 |

**Table S10.** Summary of the Wilcoxon rank sum test on sex-specific phenoloxidase activity after infection across dietary treatments

| **Treatment** | **Treatment** | **Effect** | **χ^2^** | **df** | **P** |
| --- | --- | --- | --- | --- | --- |
| Treatment A (fed on a normal diet) | Sham-infection | Sex | 1.749 | 1 | 0.186 |
|  | Infection | Sex | 0.023 | 1 | 0.881 |
| Treatment B (fed on an antibiotic-supplemented diet) | Sham-infection | Sex | 0.055 | 1 | 0.997 |
|  | Infection | Sex | 1.201 | 1 | 0.272 |
| Treatment C (fed on a microbiota-reconstituted diet) | Sham-infection | Sex | 0.365 | 1 | 0.548 |
|  | Infection | Sex | 0.142 | 1 | 0.704 |
